# *Vibrio mimicus* carrying the Type III Secretion System 2 (T3SS2) and TDH toxin of *Vibrio parahaemolyticus* in an integrative conjugative element context

**DOI:** 10.1101/2024.10.23.619894

**Authors:** Sergio Mascarenhas Morgado, Erica Lourenço da Fonseca, Ana Carolina Paulo Vicente

## Abstract

**Objectives:** Identify possible virulence factors in the genomes of *Vibrio mimicus* that may be determinants of diarrheal cases/outbreaks.

**Methods:** All *V. mimicus* genomes from Genbank were retrieved and the virulome was searched with Abricate using the VFDB database. Genomic islands (GIs) and integrative conjugative elements (ICEs) were identified using IslandViewer and ICEberg, respectively.

**Results:** Five ctx-negative *V. mimicus* genomes carrying the Type III Secretion System 2 (T3SS2) and the TDH/TRH toxin from *Vibrio parahaemolyticus* were identified. The T3SS-positive genomes presented the phylotypes T3SS2α and T3SS2β and formed two clusters and one singleton throughout the *V. mimicus* phylogeny. Genomes carrying T3SS2α and T3SS2β were associated with the *tdh* and *trh* genes, respectively. Furthermore, genomic analyses characterized an integrative conjugative element (ICE) with a size of ∼150 kb carrying both *V. parahaemolyticus* virulence determinants.

**Conclusions:** In addition to looking for cholera toxin genes in *V. mimicus* cases/outbreaks, it is important to consider *V. parahaemolyticus* toxins such as TDH and TRH. Furthermore, an ICE was identified and characterized in *V. mimicus*, being associated with T3SS2 and the TDH toxin, which is worrying, as it can disseminate these virulence factors among Vibrio spp.

## 1. Introduction

Reports of vibriosis have been increasing worldwide, mainly in Europe, Asia and the United States [1]. Among the species involved, *V. mimicus* has been associated with cases and outbreaks of gastroenteritis and cholera-like diarrhea due to ingestion of contaminated food [2]. The virulence determinants of this species are still poorly understood, since strains of *V. mimicus* that do not carry the main pathogenicity determinants of *Vibrio cholerae*, cholera toxin (CT) and co-regulated pilus toxin (TCP), could cause severe gastroenteritis [2-4]. In a recent report (2023), a severe gastroenteritis outbreak in Florida (US) was caused by *V. mimicus* strains lacking the CT and TCP, which led the authors to raise other putative candidates for the strains’ virulence, including the Type III Secretion System (T3SS) [2].

Here, analyzing the virulome of all available *V. mimicus* genomes, including those from the Florida outbreak, we detected the T3SS cluster and the thermostable direct hemolysin (*tdh*) and tdh-related hemolysin (*trh*) genes in 5/44 genomes, in regions predicted to be GIs. Most of these T3SS sequences were related to *V. parahaemolyticus* GIs. Furthermore, characterization of the T3SS genomic environment revealed that it was integrated into a region resembling an ICE, which would be the first integrative conjugative element identified in this Vibrio species.

## 2. Material and methods

All *V. mimicus* genomes from Genbank were retrieved and virulence genes were searched with Abricate (http://github.com/tseemann/abricate) using the VFDB database. GIs and ICEs were identified using IslandViewer 4 (https://www.pathogenomics.sfu.ca/islandviewer/) and ICEberg 3.0 (https://tool2-mml.sjtu.edu.cn/ICEberg3/) webservers, respectively. Gene prediction and annotation were performed with Bakta software (https://github.com/oschwengers/bakta) and manual curation.

## 3. Results and Discussion

Of the 44 *V. mimicus* genomes analyzed for virulence determinants, five presented T3SS-related genes, where 4/5 and 1/5 co-carried the *tdh* (strain N2781/GCA_008083965.1, strain F9458/GCA_009764025.1, strain E3/SRR24375803 and strain DB461B/SRR24375804) and *trh* (strain 2442/GCA_009763965.1) genes, respectively (Figure 1). While the *tdh* gene was associated to the T3SS region in the N2781 and F9458 genomes, the 2442 genome presented the T3SS and the *trh* gene in distinct genomic contexts. Therefore, our analysis revealed that for some *V. mimicus* strains that lack the main choleragenic determinants, TDH/TRH toxins would be the determining factors for gastroenteritis and/or cholera-like diarrhea.

**Figure 1.**
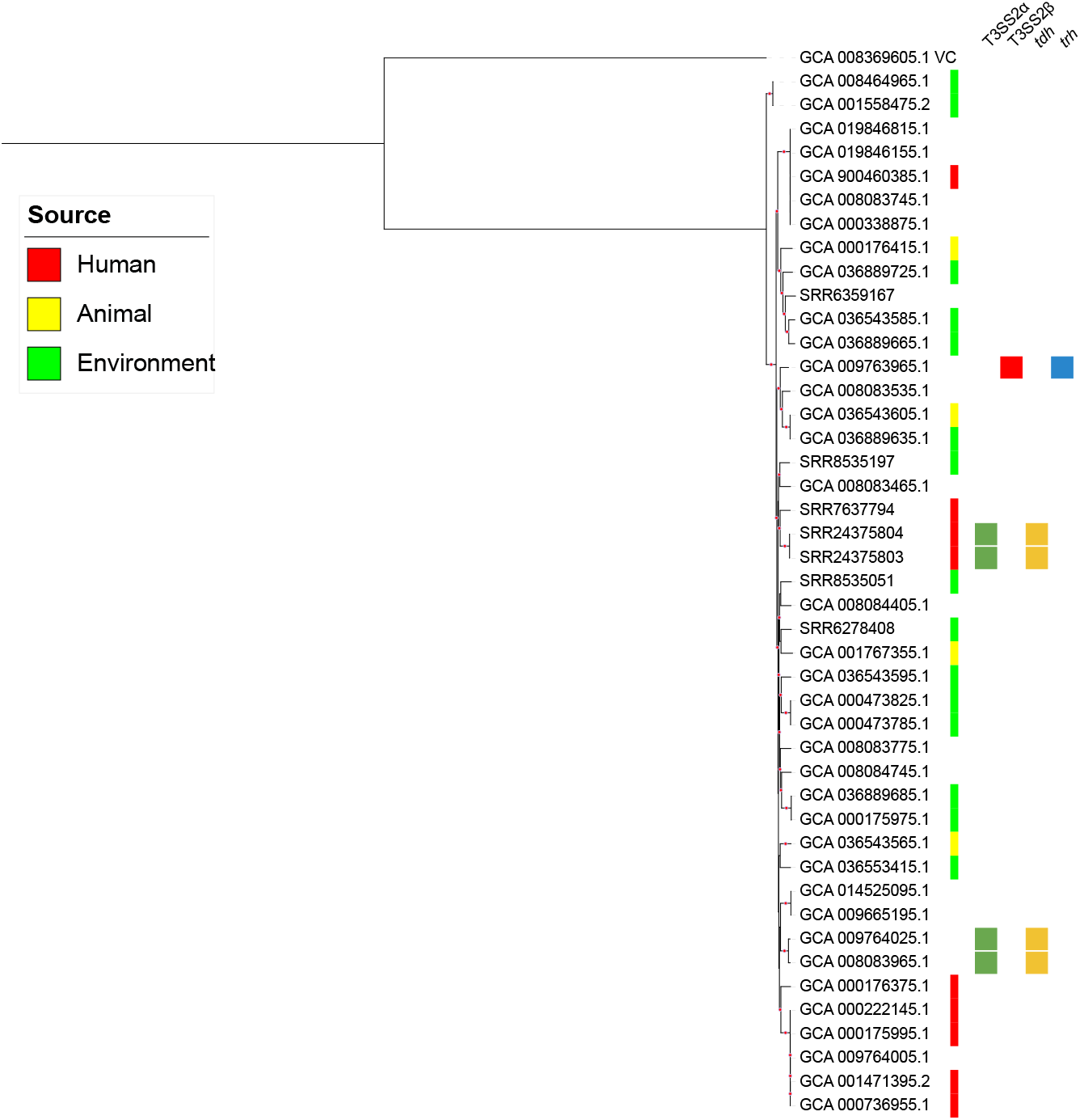
Phylogenetic analysis of *Vibrio mimicus* genomes. Maximum-likelihood tree with 1000 ultrafast bootstrap replicates built based on alignments of *V. mimicus* core genome. The best evolutionary model was GTR+F+ASC+R8, selected by the Bayesian information criterion. Colored strips represent the source of isolation. Genomes presenting T3SS2 and *tdh*/*trh* genes are associated with colored blocks. Red circles on branches represent >70% bootstrap.

In fact, the T3SS can be categorized into T3SS1 and T3SS2, where the latter is eventually associated with the *tdh* and *trh* genes. The hemolysins encoded by these genes are pore-forming toxins that damage the cell membrane and are highly prevalent in *V. parahaemolyticus* causing gastroenteritis. Indeed, these toxins have been proposed as the main virulence factors in human infections caused by this species [5]. In *V. parahaemolyticus*, these toxins and T3SS2 are part of an 80 kb pathogenicity island (PAI) (VPaI-7) [6], and similar T3SSs associated with *tdh*/*trh* genes have been reported in strains of other Vibrio species, including *V. mimicus* [6,7], although its mobility mechanism is unclear [8].

To observe whether the T3SS-carrying *V. mimicus* genomes could represent a lineage, we performed a phylogeny based on the core genome of the 44 *V. mimicus* genomes and observed that the five T3SS+ genomes formed two groups and a singleton along the phylogeny (E3 and DB461B; F9458 and N2781; 2442) (Figure 1). The *tdh* toxin gene was present in both pairs of clustered genomes and the *trh* toxin gene was in the singleton.

Furthermore, to establish the category of T3SSs carried by these *V. mimicus*, we performed a phylogenetic analysis of the T3SS based on concatenated conserved structural SctN (VscN) and SctC (VscC) amino acid sequences. This analysis showed the presence of distinct T3SS phylotypes/categories among these *V. mimicus* (Figure 2), where T3SS2α and T3SS2β were associated with the *tdh* and *trh* toxin genes, respectively (Figure 1). This association of the phylotype with the type of toxin may be biased by the number of *V. mimicus* genomes since a broader phylogeny with other species showed that both *tdh* and *trh* could be associated with different phylotypes [6]. The T3SS2α phylotype presented subgroups, where F9458 was associated with *V. cholerae* AM-19226 and SL6Y, while N2781, E3, and DB461B with *V. parahaemolyticus* RIMD2210633 (Figure 2), even though F9458 and N2781 belonged to the same lineage (Figure 1). Curiously, although the T3SS2α of F9458 was related to *V*. cholerae AM-19226, the former was associated with the *tdh* gene, flanked by an IS256 transposase, while the latter was associated with the *trh* gene, flanked by the *acf*D gene [9]. It shows the plasticity of these regions, showing variability for both the T3SS phylotype and the toxin carried among Vibrio spp.

**Figure 2.**
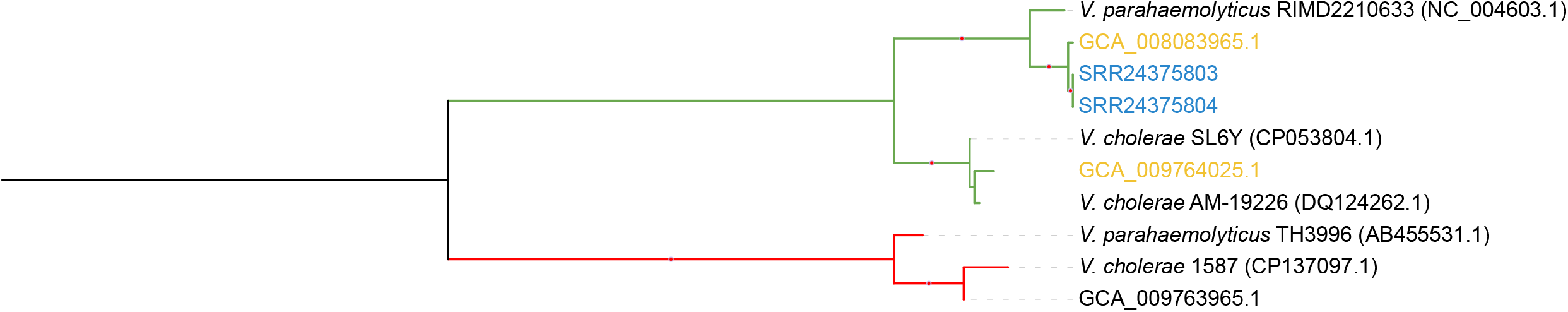
Phylogenetic analysis of *Vibrio mimicus* T3SS2. Maximum-likelihood tree with 1000 ultrafast bootstrap replicates built based on alignments of concatenated SctN and SctC amino acid sequences. The best evolutionary model was LG+G4, selected by the Bayesian information criterion. The green and red branches represent T3SS2α (associated with *tdh* gene) and T3SS2β (associated with *trh* gene), respectively. Genomes with labels of the same color belong to the same lineage. Red circles on branches represent >70% bootstrap.

Interestingly, an in-depth genome analysis of *V. mimicus* F9458 from the USA revealed that its T3SS2α gene cluster and *tdh* gene were in a region resembling an ICE, which would represent the first ICE identified in a *V. mimicus*. ICEVmF9458 was located on chromosome I and had a size of ∼150 kb and 43.4% GC content. It was possible to identify direct repeats of 12 bp (5’-TGTGTCCATTTT-3’) in the intergenic regions of both ends. The details, organization, and structure of this new ICE carrying virulence determinants common to *V. cholerae, V. parahaemolyticus*, and *V. mimicus* are in the Supplementary file. It was also possible to observe, through BLASTn analysis, that the other *V. mimicus* (N2781) from the same lineage of *V. mimicus* F9458 (USA) also harbored segments of ICEVmF9458 (Supplementary file).

In a recent *V. mimicus* outbreak in the USA (2019), the authors raised some possible virulence factors of the severe diarrhea associated with these *V. mimicus* infections, as the strains did not carry the cholera toxin genes [2]. However, analyzing the genomes of this outbreak, we were able to identify the *tdh* enterotoxin associated with T3SS2α. It has been demonstrated that T3SS2 and *tdh* are the main pathogenicity factors of *V. parahaemolyticus* strains that cause diarrheal outbreaks [7,10]. Thus, here we showed that, in addition to the previously identified T3SS, these *V. mimicus* strains also carried the *tdh* toxin gene. Both elements represent *V. parahaemolyticus* enterotoxigenic factors and, therefore, would be the determinants of this *V. mimicus* diarrhea/outbreak.

In conclusion, based on our findings, in addition to looking for cholera toxin genes in *V. mimicus* cases/outbreaks, it is important to also consider *V. parahaemolyticus* toxins such as TDH and TRH. Furthermore, we also identified and characterized the first ICE in *V. mimicus* that is associated with T3SS2 and the TDH toxin, which is concerning as it may spread these virulence factors within Vibrio spp.

## Declarations of Competing Interest

The authors have no competing interests to declare.

## Funding

This study was financed by FAPERJ - Fundação Carlos Chagas Filho de Amparo à Pesquisa do Estado do Rio de Janeiro, Processo SEI-260003/019688/2022.

## Author contributions

Ana Carolina Vicente: Conceptualization, Methodology, Writing - Original Draft, Writing - Review & Editing, Funding acquisition. Sergio Morgado: Methodology, Formal analysis, Writing - Original Draft, Writing - Review & Editing. Érica Fonseca: Writing - Review & Editing. All authors have read and approved the manuscript.

## Notes

### Competing Interest Statement

The authors have declared no competing interest.

