## Supplementary figures and images for "*Vibrio mimicus* carrying the Type III Secretion System 2 (T3SS2) and TDH toxin of *Vibrio parahaemolyticus* in an integrative conjugative element context"

### Figure S1

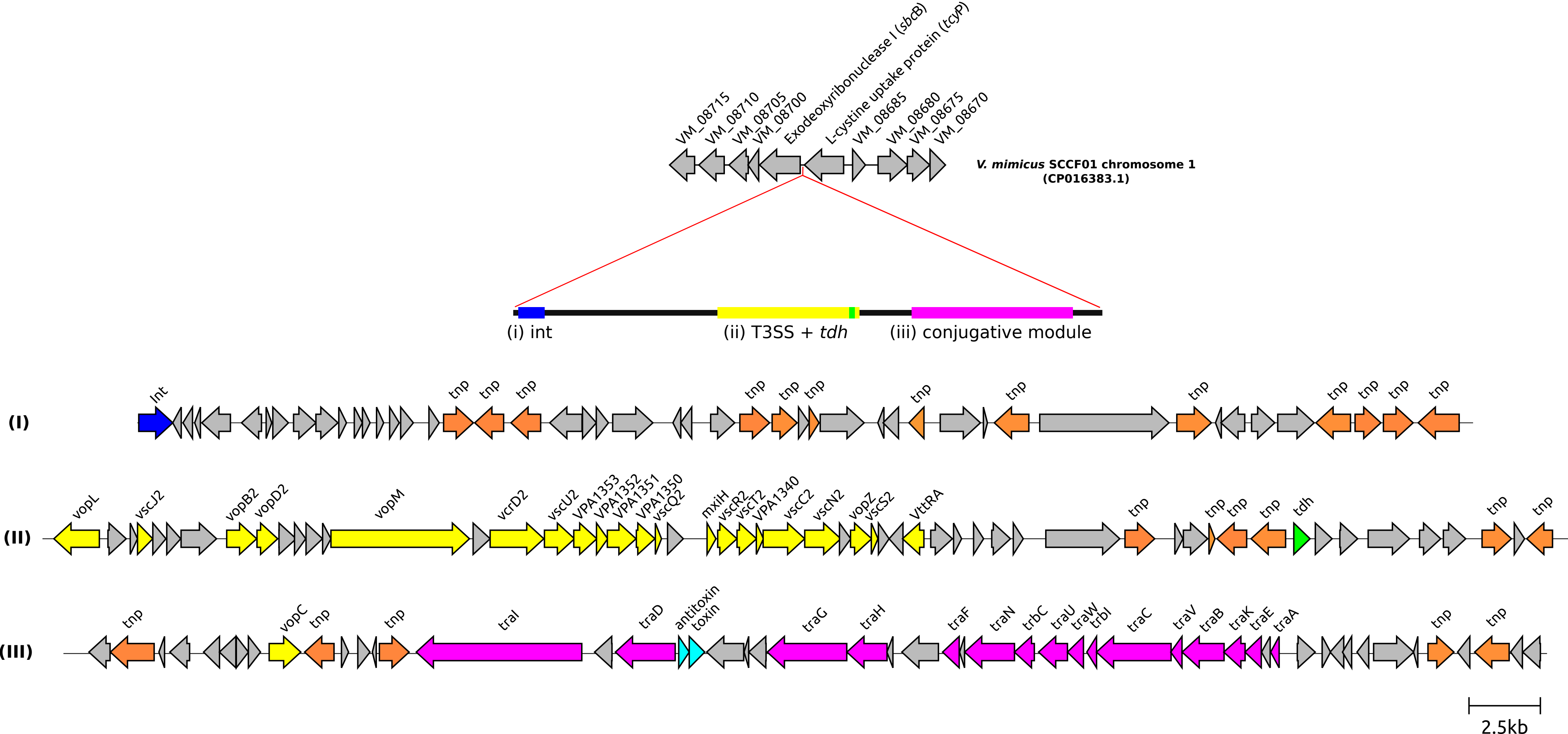
