## Supplementary File for "*Vibrio mimicus* carrying the Type III Secretion System 2 (T3SS2) and TDH toxin of *Vibrio parahaemolyticus* in an integrative conjugative element context"

**ICEVmF9458 characterization**

ICEVmF9458 was located on chromosome I of the GCA_009764025.1 genome from position 1,329,031 to 1,479,336 bp, with a size of 150,306 bp, and GC content (43.4%) similar to the genome (46.3%). It was possible to identify a direct repeat (DR) of 12 bp (5'-TGTGTCCATTTT-3') in the intergenic region of both ends. Although common in ICEs, this was not inserted into a tRNA gene [1], but between the exodeoxyribonuclease I (*sbc*B) and L-cystine transporter (*tcy*P) genes (Figure S1).

The T3SS2 region of this ICE was located at position 1,375,555 to 1,406,014 bp, including the structural (*vsc*J2/U2/Q2/R2/T2/C2/N2/S2, *vcr*D2) and effector (*vop*L/B2/D2/Z) genes. In addition, the *vtr*A (regulator), *tdh* (toxin), and *vop*C (effector) genes were in the vicinity of this T3SS (Figure S1, segment II). In contrast to *V. parahaemolyticus* T3SS2 [2], this T3SS2 was on chromosome I. Blasting this T3SS2 segment, it showed high coverage (~92-100%) and identity (~97%) with several *V. cholerae* non-01/non-o139, including the *V. cholerae* SL5Y, AM-19226, and 10432-62 genomes. Thus, corroborating with the phylogeny where this T3SS2α was more related to *V. cholerae* than to *V. parahaemolyticus* (Figure 2). But unlike the *V. cholerae* AM-19226 genome, ICEVmF9458 presented the *vsc*S gene [3].

The conjugative module of this ICE covers a region of 30,213 bp from 1,439,744 to 1,469,956 bp, being composed of *tra*I (MOBF), *tra*ABCDEFGHKLNUVW, and *trb*CI (Figure S1, segment III). Furthermore, a toxin/antitoxin system from the slvT/slvA family was observed in this module. Blastn analysis of this conjugative region only showed hits below 19% coverage with other Vibrio species. Curiously, the best hits were with *V. parahaemolyticus* plasmids (pHLD-202006, pHLD, pva2), showing 18% coverage and 73.51% identity with the *tra*I gene.

Moreover, this ICE showed a high density of transposase genes, such as IS630, IS481, IS5, IS66, IS110, IS21, IS1182 and IS256, suggesting that an ancestral GI probably acquired integrative and conjugative modules to become an ICE. In fact, the VPaI-7 lacked the integrase gene, and its region is characterized by the presence of multiple transposase genes [4].

Some similarity of this ICE was observed in the GCA_008083965.1 genome, however, it was spliced into multiple contigs. Interestingly, the conjugative region was identified with high coverage and identity (~98%) in this genome, but the T3SS region (neighboring the conjugative region) was related to *V. parahaemolyticus* (~92-97% coverage and ~97% identity). Indeed, although GCA_009764025.1 and GCA_008083965.1 belonged to the same lineage (Figure 1), the T3SSα phylogeny grouped them with sequences from *V. cholerae* and *V. parahaemolyticus*, respectively (Figure 2).

All these characteristics make ICEVmF9458 unique in relation to other ICEs in public databases, where some similarity was only observed in the T3SS2 region. Furthermore, this is the first ICE ever characterized in *V. mimicus*, while several ICEs have been described in *Vibrio cholerae* [5].

**Figure legends**

**Figure S1.** Genomic schematic of ICEVmF9458. The genetic synteny of ICEVmF9458 highlights the main regions: (i) integrase (dark blue arrow); (ii) T3SS2 gene cluster (yellow arrows) and *tdh* gene (green arrow); (III) genes related to conjugation (pink arrows). Transposase, toxin/antitoxin, and other putative genes are represented by orange, light blue, and gray arrows.
